# Catalysis by a rigid enzyme

**DOI:** 10.1101/815415

**Authors:** F. Ben Bdira, C. A. Waudby, A. N. Volkov, S. P. Schröder, E. AB, J. D. C. Codée, H.S. Overkleeft, J. M. F. G. Aerts, H. van Ingen, M. Ubbink

## Abstract

Many enzymes are dynamic entities, sampling conformational states that are relevant for catalytic activity. Crystal structures of catalytic intermediates suggest, however, that not all enzymes require structural changes for activity. The single-domain enzyme xylanase from *Bacillus circulans* (BCX) is involved in the degradation of hemicellulose. We demonstrate that BCX in solution undergoes minimal structural changes during catalysis. NMR spectroscopy results show that the rigid protein matrix provides a frame for fast substrate binding in multiple conformations, accompanied by slow, enzyme induced substrate distortion. Therefore, we propose a model in which the rigid enzyme takes advantage of substrate flexibility to induce a conformation that facilitates catalysis.

**One Sentence Summary:** The rigid matrix of BCX uses substrate flexibility in Michaelis complex formation.

## Main Text

In recent years, the combination of NMR relaxation studies and crystallography has thrown a new light on the way enzymes work (*1*). For several enzymes, it was demonstrated that their flexibility and dynamics are critical for function (*2-4*). Examples have been described in which the enzyme in one step of a catalytic cycle populates conformations as an excited state that become the ground state in the next step (*2, 5, 6*). This emerging view of the interplay between enzyme dynamics and function has recently attained much attention. Nevertheless, the generality of this model remains a matter of debate as a number of studies draw opposite conclusions (*7-12*). Enzyme conformational changes can be required to provide access to the active site (AS) to ensure specificity of the reaction or the correct order of substrate binding and product release. For enzymes with rigid folds and an accessible AS, it is unclear whether dynamics are required for function.

For example, the crystal structures of the enzyme xylanase from *Bacillus circulans* (BCX) in the resting state and various complexes are essentially identical, suggesting that no conformational changes occur during the catalytic steps [root mean square deviations (RMSD) of the C*α* atoms ≈ 0.1 Å and ≈ 0.4 Å for all heavy atoms] (**Fig. S1A**). To establish whether such changes can occur in BCX in solution, we probed the ground state structures in four states that mimic steps during catalysis, using paramagnetic NMR spectroscopy. In parallel, we determined the dynamic behavior of BCX on the millisecond timescale with relaxation dispersion (RD) NMR spectroscopy. BCX is a member of the GH11 family of glycosidases and features the conserved *β*-jelly roll fold that derives its rigidity from an extensive intramolecular hydrogen bond network involving 146 out of its 185 backbone amides, resulting in restricted dynamics on the pico-nanosecond time scale for the resting state (*13*). The shape of BCX is often compared to a right-hand fist, which includes “hand palm”, “fingers” and “thumb” regions (**Fig. 1**). The AS of the protein includes six subsites to bind units of the β1-4 xylose polymer xylan, three (–) subsites (glycon binding site) and three (+) subsites (aglycon binding site), according to the nomenclature of Davies *et al.* (*14*) (**Fig. 1C**). The glycosidic bond hydrolysis takes place between the –1/+1 subsites with a substrate binding in at least the –2/–1 and +1 subsites, in accordance with an endo-catalytic mechanism. By employing the “Koshland” double displacement mechanism, BCX hydrolyzes its substrate using a nucleophile and an acid/base catalytic dyad (*15*), proceeding through the non-covalent (Michaelis) complex (ES), covalent intermediate (EI) and product Scomplex (EP) states (**Fig. 1A**) (*16*). Crystal structures representing these states are shown in **Fig. 1C**.

**Fig. 1.**
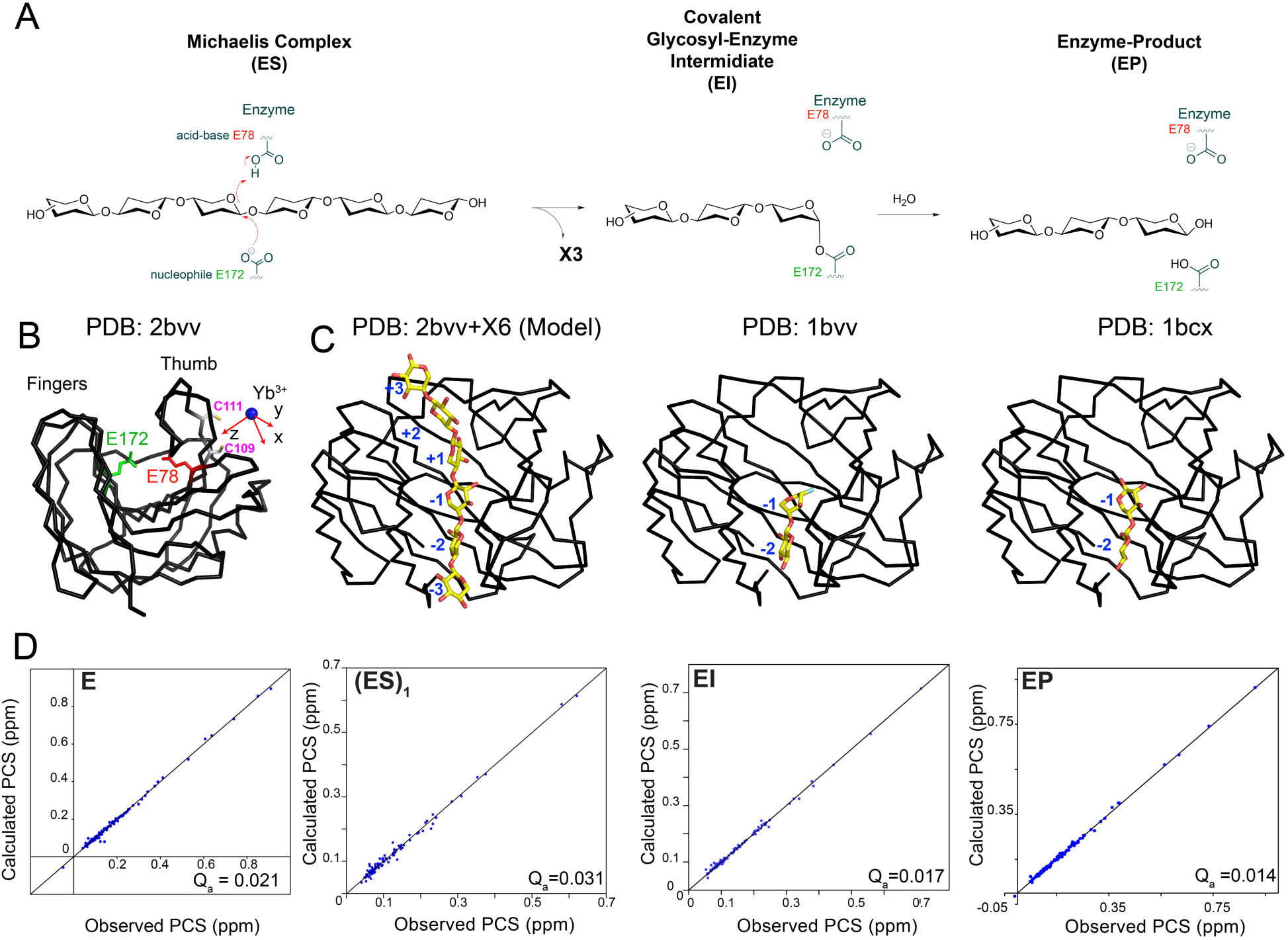
BCX structural analysis of the crystalline and solution states for several steps along the catalytic pathway. (**A**) “Koshland” double displacement retaining mechanism. (**B**) Crystal structure of resting state BCX (state E) shown as black C*α* trace (PDB:2bvv) (*20*). The nucleophile (E78, red) and acid-base (E172, green) catalytic dyad are shown in sticks. The location of the lanthanoid (blue sphere) in the CLaNP-5 tag and the frame (red arrows) defined by the anisotropic component of the magnetic susceptibility (δχ) are shown. Cysteines used for CLaNP tagging are in sticks. (**C**) Structures of BCX, from left to right, model of the ES complex based on the structure of XynII bound to X6 (PDB: 4hk8)24 in which the substrate binding subsites are numbered (*–*3/+3); EI complex (PDB: 1bvv) (*20*) and EP complex (PDB:1bcx) (*30*). The RMSD values for the Cα atoms relative to the resting state are ≈ 0.1 Å for all structures. (**D**) Correlation plots of the experimental PCS of the E, (ES)_1_, EI and EP states fitted to the crystal structure of resting state BCX (PDB: 2bvv), indicating that this structure is a good model for these catalytic states in solution. Fitting to the crystal structures of the catalytic states does not improve the fit.

### The structure of BCX in solution does not change during catalysis

Pseudocontact shifts (PCS) are a useful NMR tool to probe backbone conformation changes of proteins in solution, because PCS are highly conformation dependent (*17*). The PCS of amide nuclei of the different states were obtained with a BCX variant that has two engineered cysteine residues (T109C/T111C) linked to a lanthanoid tag (CLaNP-5) (*18, 19*), containing either a paramagnetic Yb3+ ion or a Lu3+ ion as a diamagnetic control (**Fig. 1B; S1B,C**). In the resting state, BCX shows an excellent agreement between solution and crystalline ground state structures, as revealed by the fit of the PCS to the protein crystal structure (PDB: 2bvv) (*20*) (**Fig. 1D**). The same crystal structure was also able to fit the PCS obtained for the three other states (ES, EI and EP, see below), indicating that the structural changes in the backbone of the enzyme are minimal throughout the catalytic cycle in solution, in line with the results for the crystalline states.

### Substrate and product interactions with BCX

Upon titration of a catalytically inactive variant of BCX (E78Q) with the substrate xylohexaose (X6), mimicking the Michaelis complex, a complicated chemical shift perturbation (CSP) pattern was observed. BCX is known to bind substrate in the AS, as well as in a secondary binding site (SBS) on the surface of the protein (*21*). The backbone and side chain amides of the SBS residues exhibit large CSP in the fast exchange regime (**Fig. 2A; S2A**). CSP in and around the cleft of the AS are small but widespread (**Fig. 2A; S2A**). The CSP of the AS resonances are also in the fast exchange regime and several of the amides display progressive line broadening during the titration. The remarkable difference in the amplitudes of the CSP between the two binding sites suggests a difference in the substrate binding modes. The SBS is a relatively flat region, exposed to the surface, which makes it accessible to the substrate to bind in a well-defined orientation causing large CSP (*21*). In contrast, the AS is formed by a deep cleft filled with water molecules, with six subsites. It is expected that the substrate can bind in multiple ‘registers’, i.e. shifted along the subsequent subsites and perhaps also in reverse orientation. The averaged effect of these different binding modes will be small CSP over a wide area, due to changes in the hydrogen bond network between the amides and the water molecules (*22*).

**Fig. 2.**
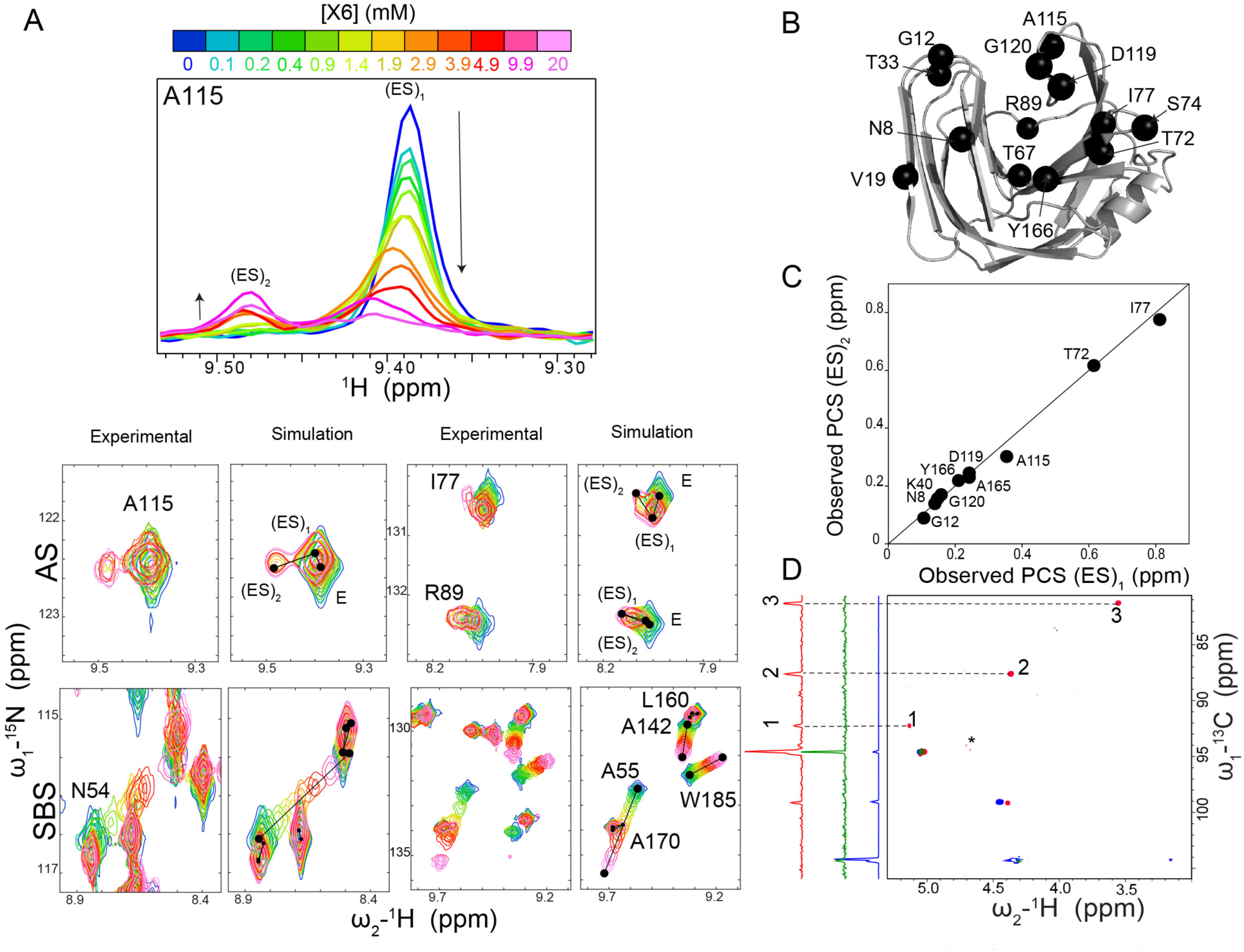
BCX Michaelis complexe formation. (**A**) In the upper panel BCX 1H-15N HSQC 1H slices are displayed for residue A115, showing that its amide resonance position and intensity changes with the appearance of its new peak in the (ES)_2_ state upon titration with X6. Note that at the highest concentrations of X6 a general line broadening occurred due to an increase in solution viscosity (**Fig. S3C**). In the lower panel, CSP of BCX E78Q backbone amides in the AS and SBS are shown next to simulated spectra using the binding model described in **M&M** and implemented in TITAN software (*31*) (see **M&M** and **Fig. S3**). (**B**) Amide groups with additional peaks representing the (ES)_2_ state shown in black spheres on BCX crystal structure. (**C**) Correlations between the experimental PCS of the (ES)_1_ and (ES)_2_ states. (**D**) Overlay between the free X6 ^1^H-13C HSQC spectrum (1500 μM in blue); in the presence of 480 μM (in green) and 1500 μM of BCX (in red). The 1D projections of the 1H-13C HSQC spectra of each condition are shown on the left side of the spectrum using the same colour code. The emerging X6 peaks are numbered and connected to the 1D projections with dashed lines. The asterisk marks the residual water signal. On the basis of the binding parameters obtained from the global fit (**Table S2**), it is estimated that the concentrations (ES)_1_ & (ES)_2_ are 74 & 44 µM and 192 & 115 µM for the samples with 480 µM and 1500 µM BCX, respectively (neglecting an allosteric effect).

Interestingly, for residues in the substrate-binding cleft of the AS a new set of resonances becomes observable at [X6] = 3.5 mM (**Fig. 2A**). This observation indicates that a second form of the enzyme-substrate complex, (ES)_2_, is present, which is in slow exchange with a first form, (ES)_1_. Although for many amides such new peaks appear, only those close to existing resonances could be assigned reliably using PCS, and these resonances are from nuclei in the binding cleft (**Fig. 2B**). At higher ligand concentrations ([X6] = 10, 20 mM) a decrease of peak intensity for (ES)_1_ resonances and increase for (ES)_2_ resonances is observed (**Fig. 2A**). A global fit of the lineshapes of the SBS and AS amides resonances (**Fig. 2A; S3)** shows that X6 binding and release from the SBS is quite fast (k_ex_ = 7.8 × 10^4^ s^-1^ at [X6] = 20 mM, k_ex_ = k_on_[X6] + k_off_). Also, the binding of X6 in the AS is in the fast regime on the NMR timescale, albeit eight-fold slower than for the SBS (k_ex_ = 0.9 × 10^4^s^-1^ at [X6] = 20 mM). The fitting model included X6 binding at two sites, the SBS and AS, as well as a slow conversion of (ES)_1_ into (ES)_2_ (**M&M** and **Table S2**). While the model fits most aspects of the titration well, it did not account for the increase of the (ES)_2_/(ES)_1_ ratio at the high concentrations of X6. This increase indicates that the (ES)_2_ state, being the state with an X6 molecule in the AS in conformation 2, becomes more populated when a second X6 molecule occupies the SBS, suggesting an allosteric effect of X6 binding in the remote SBS on the AS. Inclusion of such effect in the modelling led to overfitting and, thus it could not be modelled reliably. Interestingly, it has been reported before that the longer substrates xylododecaose (X12), as well as soluble and insoluble xylan, can bind to the AS and SBS simultaneously with a single substrate molecule, enhancing the K_m_ of the enzyme (*21*). Thus, information transfer between the SBS and AS sites appears to be possible. The (ES)_1_ and (ES)_2_ states exchange at a rate of 71 s^-1^ (k_ex_ = k_forward_ + k_backward_). This exchange rate is in the range of the turn-over rate of BCX using the artificial substrate PNP-X2 (20 s-1, 23°C), for which the glycosylation step has been shown to be rate limiting (*23*). Thus, these observations suggest that the conversion of (ES)_1_ into (ES)_2_ could be related to the rate-limiting step in catalysis. A similar slow transition process was reported for a different type of glycosidase, chitosanase, when titrated with chitosan hexaose (*24*). In that study, the slow exchange was attributed to a conformational change of the protein induced by ligand binding. However, for BCX the similarity of the PCS of resonances in (ES)_1_ and (ES)_2_ indicates that the conversion from (ES)_1_ to (ES)_2_ does not lead to a major structural rearrangement of the enzyme (**Fig. 2C**).

This observation poses the question what the slow transition step in BCX Michaelis complex represents. We wondered if a conformational change in the bound substrate could be the cause of the two ES states. To test this hypothesis, we observed the effects of BCX binding on X6 by using 1H-13C HSQC NMR experiments at natural abundance to detect nuclei of the sugar rings. Many of the 13C resonances lie in a region of the spectrum (75-105 ppm) in which the protein shows no resonances. Upon addition of BCX, the resonances of many X6 resonances shift and broaden beyond detection, even at a BCX:X6 ratio of 0.2:1 (**Fig. S2D**), in line with the model formulated above, in which X6 molecules bind and dissociate rapidly in different modes and thus different chemical shifts, both to the SBS and the AS. Then, the ratio of BCX and X6 was increased to 1:1 (1.5 mM each). Interestingly, several new peaks were detected in the region of the sugar resonances, indicating the presence of another form of X6 that is not affected by the fast-exchange binding events (**Fig. 2D**). Thus, this observation is analogous to the observation of the additional peaks observed for the BCX amides, the (ES)_2_ state. The ^13^C chemical shifts of the new peaks differ considerably (> 10 ppm) from those of the free sugar, indicating that the new form must experience significant conformational change, because carbon chemical shifts are dominated by conformational effects. A crystal structure of X6 bound to another GH11 xylanase, XynII (PDB: 4hk8) shows that the oligosaccharide chain is in a well-defined, yet strained conformation (*25*). This structure could represent the equivalent of the BCX (ES)_2_ structure. The authors suggest that this state could be primed for catalysis. A computational study supported the observation that the substrate is distorted, but it suggested that the observed conformation of the sugar is off the predicted trajectory for catalytic conversion (*26*).

To mimic the next step in the catalytic cycle, the covalent intermediate state, a covalent inhibitor (epoxyX2) was used that resembles xylobiose (X2) (*27*). Despite the absence of structural changes in the enzyme (see above), adduct formation leads to many CSP (**Fig. S2B**), which are attributed to changes in the hydrogen bond network in and around the AS due to binding of the adduct and displacement of water molecules. The last step of the reaction is product release from the glycon pocket after hydrolysis. A NMR titration of BCX with X2, representing the product, shows that ligand binding is in the fast exchange regime, with k_ex_ = 4.3 × 10^4^s^-1^ at [X2] = 130 mM (**Fig. S2E**). No additional peaks in the slow exchange regime were observed at high product concentration, contrary to the titration with X6. X2 also does not interact with the SBS to a detectable level. Based on the CSP pattern, the ligand appears to occupy only the -1/-2 subsites (**Fig. S2C**), indicative of a difference in the binding sites affinities between the glycon and aglycon sites toward the product.

### Substrate and product binding enhances exchange effects

RD-NMR experiments were performed to detect lowly populated states that are in dynamic equilibrium with the ground states. RD-NMR on the resting state of BCX show that millisecond chemical exchange is limited to residues located within the fingers and the thumb regions, with few residues affected in the AS (**Fig. S4A; S5A**). A two-site exchange model fit of the dispersion data yields a k_ex_= 2.4 (±0.1) × 10^3^ s^-1^ (20°C) and a population of the excited state p_B_ of 0.8% (**Table S3**). It is noted that a two-site exchange model is the simplest case and it may represent an oversimplification (*28*). The analysis is used here to estimate the order of magnitude of the k_ex_ and p_B_. For the EI complex, the chemical exchange effects mostly resemble those for the resting state BCX, both qualitatively and quantitatively (k_ex_ = 2.05 (± 0.08) × 10^3^ s^-1^, p_B_ of 0.6%, **Fig. S4C; S5B** and **Table S3**). Large exchange broadening effects (R_ex_ up to 50 s^-1^) were observed for both the Michaelis complex (ES)_1_ and the product complex (EP) for amino acid residues that form the binding cleft (**Fig. 3**). A two-site exchange fit yielded k_ex_ = 741 (±9) s^-1^, p_B_ = 9.2 (±0.2) % for ES_1_ and k_ex_=1.53 (±0.02) × 10^3^ s^-1^, p_B_ = 4 (±0.04) % for EP (**Fig. S4B, D; Table S3**;). In the EP complex, large R_ex_ values were found in particular for amide nitrogen atoms of the thumb and fingers at the -1/-2 subsites, which also experience CSP upon ligand titration (**Fig. S2C**).

**Fig. 3.**
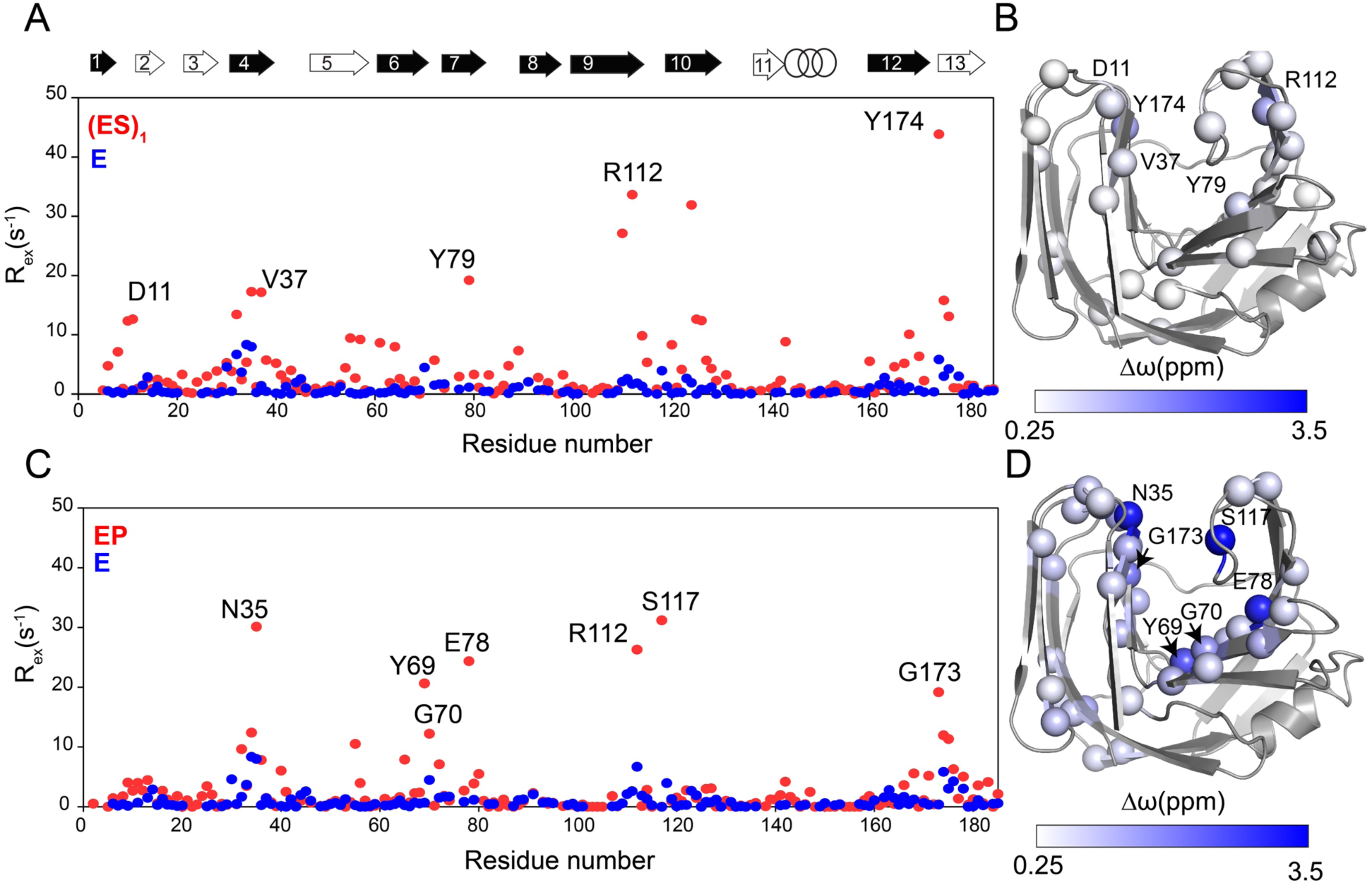
Non-covalent interactions enhance BCX millisecond time scale dynamics in the (ES)_1_ and EP states. Overlay of the R_ex_ values of (ES)_1_ (**A**) or EP (**C**) (red dots) and E states (blue dots) plotted versus the residue number. Several residues that show a prominent difference in R_ex_ are labelled. The secondary structure elements of BCX are represented by black arrows for β-strands of sheet A and in white ones for sheet B and the α-helix in rings. The “thumb” loop connects β-strands 9 and 10 (*21*). (**B**) and (**D**) Amide nitrogens used in RD global fit are shown in spheres on the BCX crystal structure (PDB ID: 2bvv) colored by their δω between the ground and excited states, using a white/blue gradient. The δω were derived from a global two-state fit to the dispersion curves.

### Origin of chemical exchange effects in the non-covalent complexes

Crystal structures of BCX in various states as well as the PCS analysis indicate that BCX shows negligible conformational changes during catalysis, raising the question what conformations the excited states observed in the RD NMR experiments represent and whether they are relevant for catalysis. For other enzymes, the evidence that excited states represent conformations of the next state in the catalytic cycle has been obtained by showing a correlation between the 15N chemical shifts of excited states, derived from the δω values of RD NMR experiments, with those of the successive ground states (*2, 5, 6*). Such a correlation could not be found for the data reported here (**Fig. S6**), in line with a similar analysis on a BCX related protein, XlnB2 (*29*). The combination of a lack of enzyme conformational changes and the strong enhancement of the chemical exchange for the states in which the enzyme interacts non-covalently with substrate or product molecules ((ES)_1_ and EP states), suggests the presence of multiple binding modes of the substrate in the AS. As mentioned above, it can be expected that both X6 and X2 can bind in different ‘registers’ and positions of the six binding subsites, with different affinities. Each binding mode would change the hydrogen bond network of amides and waters in the AS in a different way and, thus, cause different CSP. The presence of multiple binding modes in (ES)_1_ can explain the observed chemical exchange in the AS. The NMR titration results for (ES)_1_ formation indicated rapid association and dissociation (k_ex_ ≈ 10^4^ s^-1^), which would be too fast to yield RD effects. However, the exchange between bound states *via* the free state can still be sufficiently slow to cause chemical exchange broadening, as shown in the **M&M** and **Table S4** for a simple case with two binding modes, (ES)_1_ and (ES)_1_^*^. The fact that in the resting enzyme the affected amides are in loop regions outside the AS in conjunction to the very low population of its excited states suggests that the enzyme dynamics may not be relevant for substrate binding or product release.

All the NMR results can be integrated in a model describing the successive states of the catalytic cycle, **Figure 4**. The titration results indicate rapid association of BCX and X6 into the non-covalent complex (ES)_1_, in the enzyme binding cleft. The substrate can bind in the multiple modes, resulting in different (ES)_1_ complexes, causing the observed RD effects for the amides in the AS. One or more of (ES)_1_ complexes can proceed to form (ES)_2_, with a high-energy transition barrier, because the conversion is slow. This conversion is attributed to the substrate distortion in the AS without inducing conformational rearrangement in the protein, an induced fit of the substrate. The rate of conversion between (ES)_1_ and (ES)_2_ is in the range of the rate-limiting glycosylation step (*23*), so (ES)_2_ may well represent an activated form of the ES complex that can proceed to form the EI complex. After hydrolysis, the product complex (EP) is in fast exchange with the free enzyme and product, as shown by the NMR titration, and, analogous to the (ES)_1_ complex, the product can bind in various modes, resulting in multiple EP complexes, as evidenced by the RD effects for this complex.

**Fig. 4.**
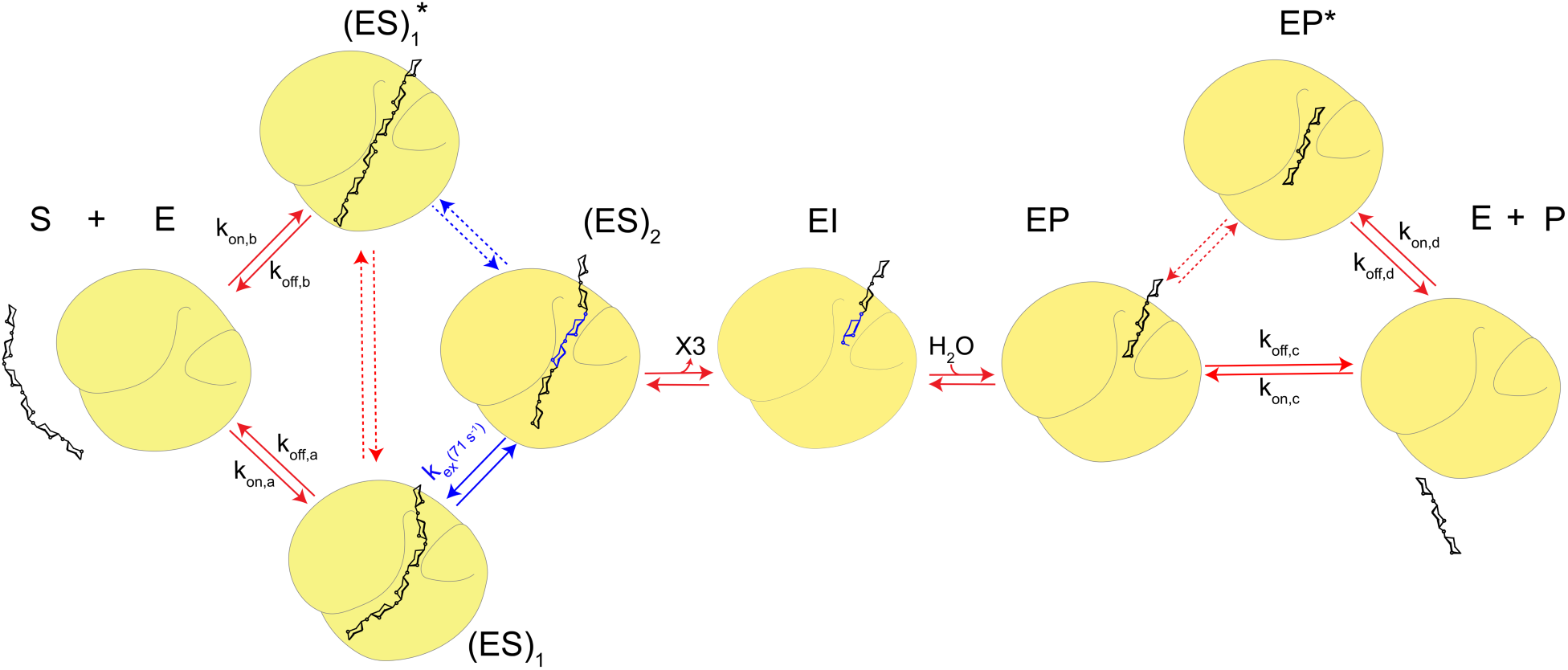
Proposed model for BCX catalytic cycle. Blue sugar units indicate X6 distortion. Possible values for the microscopic rates k_on,i_ and k_off,i_ that yield the experimental off rates, exchange rates, minor state populations and dissociation constants are given in **Table S4**. Dashed arrows indicate equilibria that could be present but for which no experimental evidence was obtained. In the model, the substrate X6 and X2 can bind in multiple ways, forming a major state and one or more minor state(s), indicated with an asterisk.

In conclusion, xylanase from *B. circulans* is an enzyme with remarkably few conformational changes during catalysis. It is concluded that the X6 substrate can bind in multiple modes with different affinities and high dissociation rates. A slow transition occurs upon binding of the substrate X6, which we ascribe to substrate distortion, and that could represent a rate-limiting event in the catalytic cycle. Contrary to the recent insights for other enzymes, the enzyme appears to provide a rigid frame that stabilizes specific substrate conformations to enable the reaction to proceed. The observed dynamics in the Michaelis complex is attributed to multiple substrate binding modes within the AS. Dynamics in AS generally reduces protein stability 29, thus, a design of a rigid enzyme that is able to take advantage of the inherent conformational landscape of the substrate, may offer evolutionary advantages. This study accentuates the importance of studying not only enzyme but also substrate dynamics to obtain a complete picture of enzyme catalyzed reactions.

## Acknowledgments

We thank Professor Lewis E. Kay for providing us with the TROSY-CPMG pulse program.

## Author contributions

F.B.B. performed the experiments. F.B.B.; C.W.; A.V.; E.B; H.V.I and M.U analyze the data. S. P. S.; J. D. C. C.; H.S. O., J. M. F. G. A. provided the tools. F.B.B. and M.U. supervised the project; F.B.B. and M.U. wrote the manuscript. All authors reviewed and corrected the manuscript.

## Supplementary Materials

Materials and Methods

Figures S1-S8

Tables S1-S4

References (*1-12*)

## Supplementary Materials for

### Materials and Methods

#### Materials

The epoxyX2 was synthesized as reported by Schröder *et al.*^1^. CLaNP-5 was synthesized and loaded with lanthanoids as published ^2,3^. Xylobiose (X2) and Xylohexaose (X6) were purchased from Megazyme and used without further treatment.

#### Methods

##### Mutagenesis

BCX E78Q, BCX T109C/T111C and BCX E78Q T109/T111C mutants were prepared by site-directed mutagenesis using the Quik-Change protocol. The presence of the mutations was confirmed by sequencing.

##### BCX production and purification

The gene of BCX with a sequence optimized for expression in *Escherichia coli* was purchased from Life Technologies, including a N-terminus 6-His-Tag followed by a TEV protease recognition site. The gene was subcloned into plasmid pET28a using NcoI/XhoI restriction sites. *E. coli* BL21DE Rosetta cells were transformed with the obtained vector and cultured in 2 mL lysogeny broth containing 100 μg/mL kanamycin for 6 hours at 37°C. Of the miniprep culture, 100 µL were used to inoculate 50 mL minimum medium supplemented with ^15^N labeled ammonium chloride (0.3 g/L) and either normal glucose (4 g/L) or uniformly labelled ^13^C-glucose (2 g/L). The next day, 0.5 L of the same medium was inoculated with the overnight pre-inoculation culture and incubated for 8 hours at 37°C with shaking until the OD_600_ reached 0.6. BCX gene expression was induced with 0.5 mM IPTG and the culture was incubated overnight at 20°C with shaking. The cells were harvested by centrifugation and the pellet was resuspended in an extraction buffer (50 mM phosphate buffer solution (PBS), 40 mM imidazole, pH 7.5) and sonicated. The resulting 6His-BCX supernatant was purified by Ni^2+^ affinity chromatography. Subsequently, the eluted BCX fractions were collected and the protein was brought into a solution of 50 mM PBS using a PD10 column. The 6-His-BCX protein was incubated with TEV protease overnight at 4°C. Next, the cleaved 6-His-Tag and the TEV protease were separated from BCX using Ni^2+^ affinity chromatography. The flow-through fraction, which included BCX, was collected and the buffer was changed to 25 mM sodium acetate, pH= 5.8, using a PD10 column. The protein was concentrated with Amicon® Ultra-Centrifugal Filters with 10 kDa cutoff and the purity was checked by SDS-PAGE (estimated to be >95%). The BCX concentration was determined by UV absorbance, using the theoretical extinction coefficient of 81790 M^-1^cm^-1^ (PROTEIN CALCULATOR v3.4: http://protcalc.sourceforge.net).

#### EI covalent complex formation

The protein was incubated with 20 molar equivalents of epoxyX2 at 30 °C (**Fig. S7A**). The reduction of the enzyme activity was monitored over the course of incubation using 4MU-xylobiose as a substrate (Megazyme) (**Fig. S7B)**. The reaction rates were monitored by measuring the fluorescence of the released 4MU at its emission wavelength of 445 nm on a Cary Eclipse Fluorescence Spectrophotometer (Agilent). After an overnight incubation, the excess of epoxyX2 was removed using a PD10 column and complex formation was checked by electrospray mass spectrometry (Waters). The HSQC spectrum of the EI was recorded and compared to the one of the resting state protein to check adduct formation (**Fig. S7C**). The same procedure was applied to prepare a ^15^N, ^13^C BCX-epoxyX2 sample (for assignment), ^15^N BCX-epoxyX2 sample (for ^15^N CPMG RD NMR) and ^15^N BCX T109C/T111 CLaNP-5-(Lu^3+^/Yb^3+^)-epoxyX2 samples for PCSs measurements.

#### NMR spectroscopy

Sequential assignment of the BCX WT at 20 °C was performed by using triple resonance HNCACB and CBCAcoNH spectra on a ^15^N, ^13^C BCX NMR sample (0.6 mM in 25 mM sodium acetate, pH= 5.8) and in combination with the published data (BMRB entry code: 4704) ^4^. The obtained peak list was used to assign the BCX E78Q, BCX T109C/111C and BCX E78Q T109C/111C mutants backbone amides. The assignments of these were checked and corrected using PCS (**Fig. S8A**). The same assignment procedure as for the BCX WT was applied for the assignment of the BCX-epoxyX2 (EI complex). In the EI spectrum the amide resonances of three residues were missing, presumably due to peak broadening, including E78, which is supposed to covalently bind to the C1 of the ligand, G70 and S117, which are in contact with the -1 subsite xylobiose sugar unit, according to BCX-2FXb crystal structure (PDB ID: 1bvv) ^5^. The NMR spectra were acquired on Bruker Avance III (HD) 600 or 850 MHz spectrometers, equipped with TCI cryoprobes, processed by TopSpin 3.5 (Bruker Biospin) and analysed with Sparky software ^6^.

#### NMR titrations and peak shape analysis

The NMR titration of ^15^N BCX E78Q (123 μM) in 25 mM sodium acetate (pH=5.8) and 6% D_2_O with an increasing X6 concentration (123 μM - 20 mM in the same buffer) was performed at 20 °C. Some general line broadening was observed (**Fig. S3C**), due to viscosity increase induced by X6. The same procedure was applied when titrating BCX wild type with X2 (1.3-130 mM). The ^1^H-^15^N HSQC spectrum of every titration point was recorded and the average chemical shift perturbation values (Δδ_avg_) relative to the free protein were calculated using the last point of the titration (in the presence of 130 mM X2 and 20 mM X6) according to the equation (1):

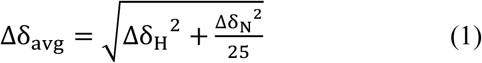

Two-dimensional lineshape fitting was performed in MATLAB (R2018a, TheMathWorks) using TITAN (v1.6) ^7^ with a customized binding model (see the scheme below). State **A** represents the resting state enzyme; state **B** is the enzyme with substrate bound to the AS in fast exchange, with the SBS free; state **C** is the state with distorted substrate in the AS with the SBS free; state **X, Y** and **Z** represent the equivalent states as **A, B** and **C**, respectively, with the SBS occupied by X6. States **B**+**Y** represent (ES)_1_, states **C**+**Z** (ES)_2_. The K_D_, k_off_ and k_ex_ values with the same color in the scheme were constrained during the fitting to be identical. 19 residues were fitted from the BCXE78Q titration with X6, representing AS and SBS residues with a variety of maximal chemical shift differences. Peak positions and linewidths of free protein amide resonances were first fitted and then held fixed during successive rounds of fitting. Additional constraints were imposed to reduce the number of free parameters: A single linewidth was fitted for all ligand-bound states of a resonance; the chemical shifts of AS/SBS site resonances as well as the dissociation or exchange rates were assumed to be independent of the absence or presence of X6 at the other site. These constraints almost completely eliminate the possibility of fitting an allosteric effect of SBS binding on the binding or conformational change in the AS. The obtained values are listed in **Table S2**. Reported parameter uncertainties were determined using a new jackknife resampling algorithm, which yielded more conservative error estimates than the block bootstrap resampling approach (*7*) (**Fig. S3A,B**). The BCX wild type titration with X2 was fitted to a two-state binding model (**Fig. S2E**).

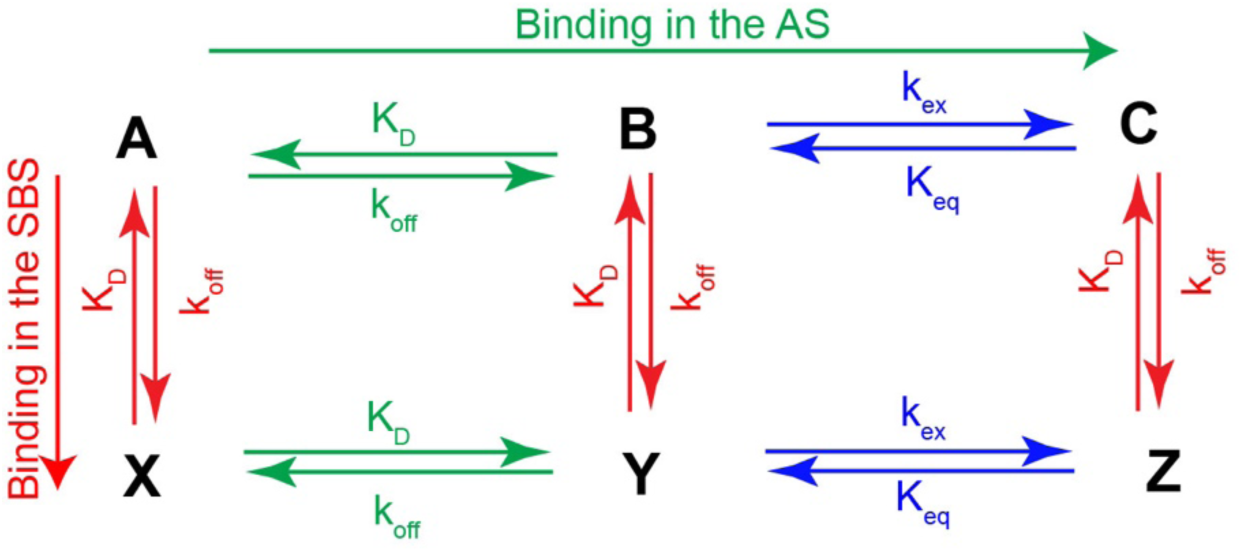

#### Paramagnetic NMR spectroscopy

Uniformly labelled ^15^N BCX T109C/T111C was incubated with 10 mM DTT in 50 mM PBS (pH=7.2) and 150 mM NaCl for 1 h on ice to reduce possible disulphide bridges. DTT was removed by using a PD10 column. The protein was mixed with 10 molar equivalents of CLaNP-5-Lu^3+^ (diamagnetic sample) or CLaNP-5-Yb^3+^ (paramagnetic sample) and left stirring for 2 h at room temperature. The samples were concentrated to 0.5 mL using a 10 kDa Amicon® Ultra-Centrifugal Filter and injected into a Superdex G75 analytical gel filtration column to remove unreacted tag and protein dimers. To verify the labelling efficiency, the protein samples were analyzed on 15 % SDS-PAGE in the presence and absence of a reducing reagent (*β*-mercaptoethanol) and checked by acquiring ^1^H-^15^N HSQC spectra. Weak peaks of untagged BCX were observed for both samples (<5% unlabeled). To obtain the experimental PCSs, the frequency differences between the analogous resonances in the paramagnetic and diamagnetic samples were calculated. For each state, the experimental PCS were fitted to the crystal structure of the resting state (PDB ID: 2bvv) ^5^ using Numbat software ^8^, yielding the location of the lanthanoid relative to the protein and the δχ tensor size and orientation. The lanthanoid position was ∼ 8-10 Å from the C*α* atoms of the engineered cysteine residues. The jackknife error analysis implemented in Numbat was used to estimate the errors (**Fig. S1B**). Only very subtle changes in the tensor positions were observed between the different states (**Fig. S1A**). The Q_a_ factor of the experimental PCSs fit to the BCX crystal structure was calculated using the equation (2) ^9^:

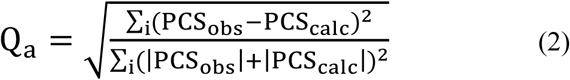

Where i sums over all PCSs. Note that the Q_a_ differs from the regular Q by addition of the observed and calculated PCS in the denominator. This definition makes the factor unbiased toward those points for which PCS_calc_ > PCS_obs_, as compared to points for which PCS_obs_ > PCS_calc_.

#### NMR relaxation dispersion and CPMG data analysis

^15^N CPMG RD data sets were recorded at 20°C using a ^15^N TROSY-CPMG pulse scheme ^10^ on a 0.5 mM BCX sample in 25 mM sodium acetate (pH=5.8), 6% D2O. Data sets were recorded at field strengths of 14.2 T and at 19.9 T. The constant time relaxation delay, T_relax_, was set to 50 ms. Dispersion profiles comprised ∼ 20 different ν_CPMG_ frequencies, recorded in an interleaved manner, with values ranging from the minimum possible value (1/T_relax_) to a maximum of 1000 Hz. Errors were estimated on the basis of repeat measurements at three different *v*_CPMG_ frequencies. Each CPMG dataset required approximately 14 hours of measurement time. All experiments were performed on Bruker spectrometers equipped with a TCI-Z-GRAD triple resonance cryoprobe. The peak shape fit of the 3D-pseudo planes was performed using FuDa ^11^. The R_2,eff_ (ν_CPMG_) were obtained via the relation (3):

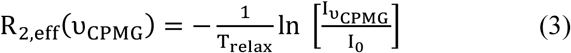

Where I(ν_CPMG_) and I_0_ are peak intensities with and without the 50 ms delay, respectively. Values of the exchange parameters were extracted from a global fit of R_2,eff_ (ν_CPMG_) profiles to a two-site exchange model using CATIA (http://www.biochem.ucl.ac.uk/hansen/catia). R_ex_ values were calculated using the formula R_ex_= R_2,eff (20 Hz)_ - R_2,eff (1000 Hz)_ from the experimental data. No major effects of the E78Q mutation on the BCX ms chemical exchange were observed, as can be concluded from the overlay of the experimental R_ex_ values of wild type enzyme and the BCX E78Q variant (**Fig. 8B**).

#### Modelling of kinetic parameters

To explain RD effects due to a ligand binding in two or more modes to the enzyme, a model is described in which we consider the conversion of (ES)_1_^a^ to (ES)_1_^b^ via free E (eq. 4). The k_ex_ for conversion (ES)_1_^a^ into (ES)_1_^b^ and back is given by eq. 5.

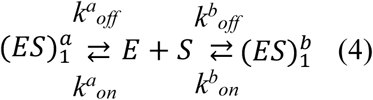

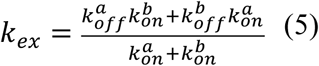

---

**Fig. S1.**
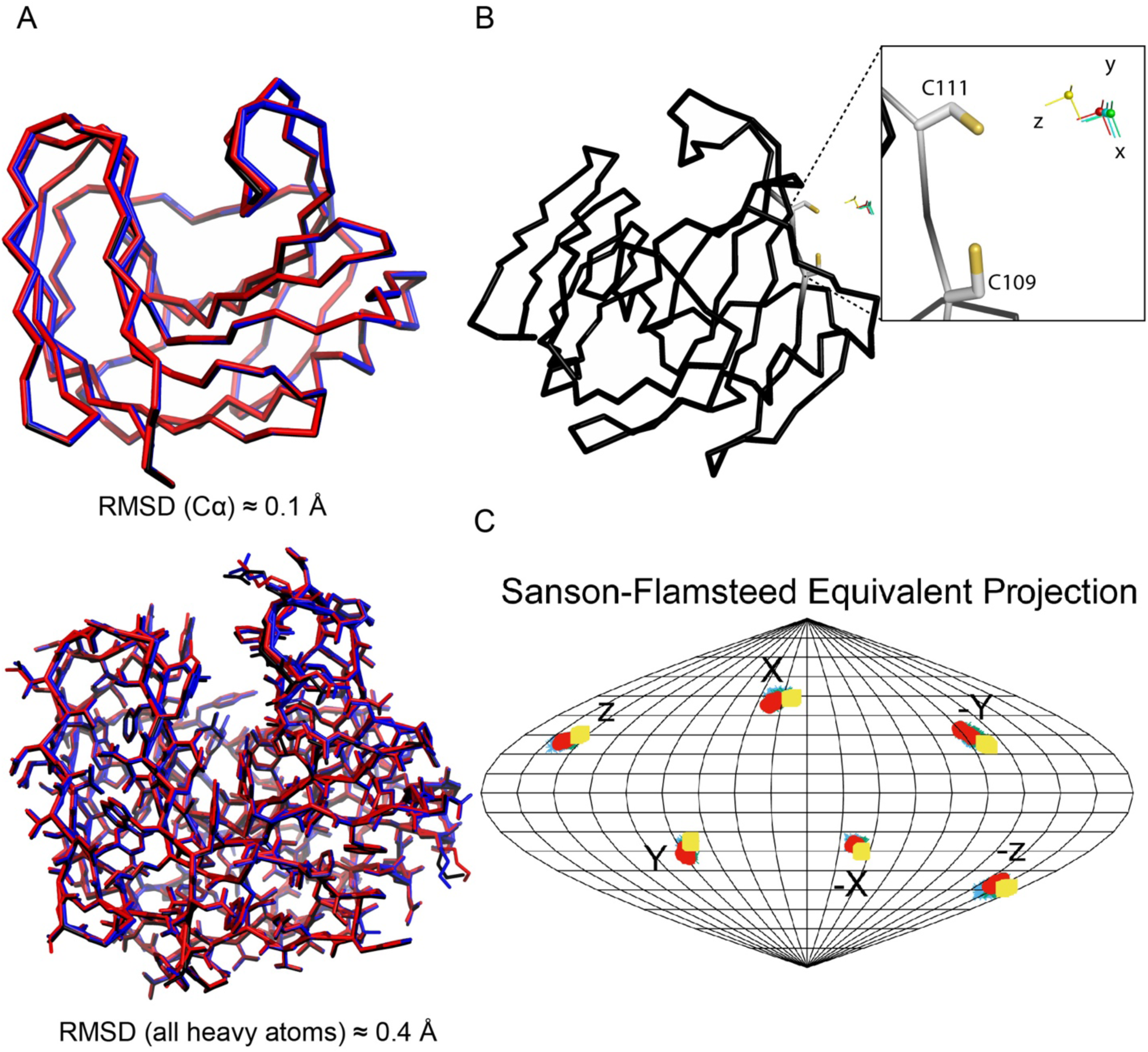
BCX in the E, (ES)_1_, EI and EP states crystal structures and tensors parameters. (**A**) Overlay of BCX crystal structures in the free (PDB: 2bvv, black ribbon), EI (PDB: 1bvv, blue ribbon) and EP (PDB:1bcx, red ribbon) states. Upper and lower panels are for the overlay between the structures C*α* atoms and all heavy atoms, respectively. (**B**) Tensors positions of the BCX free (cyan); (ES)_1_ (yellow); EI (red) and EP (green) states plotted on the protein crystal structure (PDB: 2bvv) obtained after fitting to the experimental PCS obtained for each state. (**C**) One hundred δχ tensors generated from jackknife analysis within Numbat for BCX tensors using the same color code as above. The locations where the X, Y and Z components pierce the sphere are indicated. The surface of the sphere is shown as a Sanson-Flamsteed equivalent projection with gridlines at every 20° (vertical) and 10° (horizontal). The obtained tensor parameters are listed in **Table S1**.

**Fig. S2.**
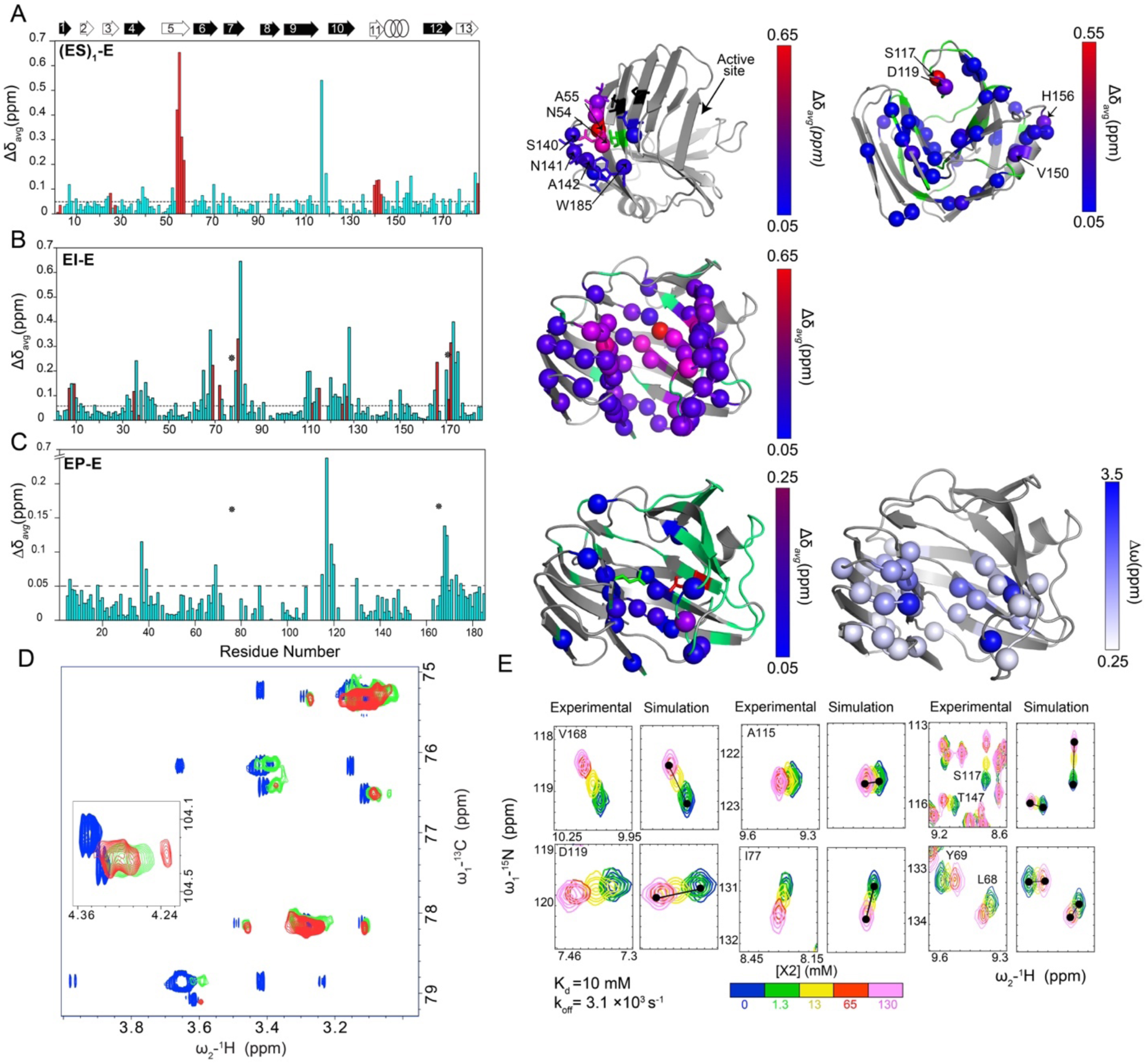
BCX binding sites in the (ES)_1_, EI and EP states. (**A**) Left panel shows the average CSP (*Δ*δ_avg_) between the (ES)_1_ and Estates. The *Δ*δ_avg_ were obtained from the highest point of the titration in the presence of 20 mM X6 (>85% bound state). *Δ*δ_avg_ of the SBS amides are shown in red bars as determined in (*4*). Middle panel represents the mapping of the SBS amide groups with *Δ*δ_avg_ > 0.05 on the BCX crystal structure (PDB ID: 2bvv) ^5^ in spheres and sticks and colored by *Δ*δ_avg_ using a blue/red gradient. Residues for which no data were obtained are colored in green and the ones with *Δ*δ_avg_< 0.05 ppm in black. Right panel shows the amide groups of the AS with *Δ*δ_avg_ > 0.05 ppm mapped on the BCX crystal structure (PDB ID: 2bvv) in spheres and coloured by *Δ*δ_avg_ using a blue/red colour gradient. Residues for which no data were obtained are coloured in green and the ones with *Δ*δ_avg_< 0.05 in grey. (**B**) Left panel shows the *Δ*δ_avg_ for the CSP between EI and E states obtained are plotted versus BCX residue number. Residues that form direct interactions with the ligand according to BCX crystal structure (PDB: 1bvv) are shown in red bars. The positions of the E78/E172 catalytic dyad are indicated with asterisks. The color coding is the same as in (**A**). Right panel is the mapping of the *Δ*δ_avg_ on the BCX crystal structure. (**C**) Left panel represents the *Δ*δ_avg_ between the EP and E states plotted against BCX residues numbers in the presence of 130 mM X2 (>90% bound state). Positions of the E78/E172 catalytic dyad are indicated with asterisks. The color coding is the same as in (**A**). Right panel indicates the amide nitrogens with enhanced millisecond dynamics in the EP state mapped on the BCX crystal structure (PDB ID: 2bvv). The residues are colored by their δω between the ground and excited states, derived from a global two-state fit to their dispersion curves, using a white/blue gradient. (**D**) X6 conformational states in the BCX active site. X6 ^1^H-^13^C HSQC spectra of the titration with BCX are overlaid, showing the region of X6 signals that does not overlap with protein signals. Blue, green and red spectra are for the free X6 (1500 μM); in the presence of 300 μM and 480 μM of BCX, respectively. The insert represents the chemical shift perturbation of the xylose units’ anomeric carbons using the same colour code. Note the extensive exchange broadening of the sugar signals. The samples were prepared in 10 mM deuterated sodium acetate buffer, pH 5.6 in 100% D_2_O and the measurement was performed at 20 °C. (**E**) Line shape analysis of the BCX titration with X2 (EP state) using two state binding model implemented in TITAN. Binding parameters obtained from line shape analysis are K_d_ = 10.0 (±0.1) mM and a k_off_ = 3.1 (0.2) × 10^3^ s^-1^

**Fig. S3.**
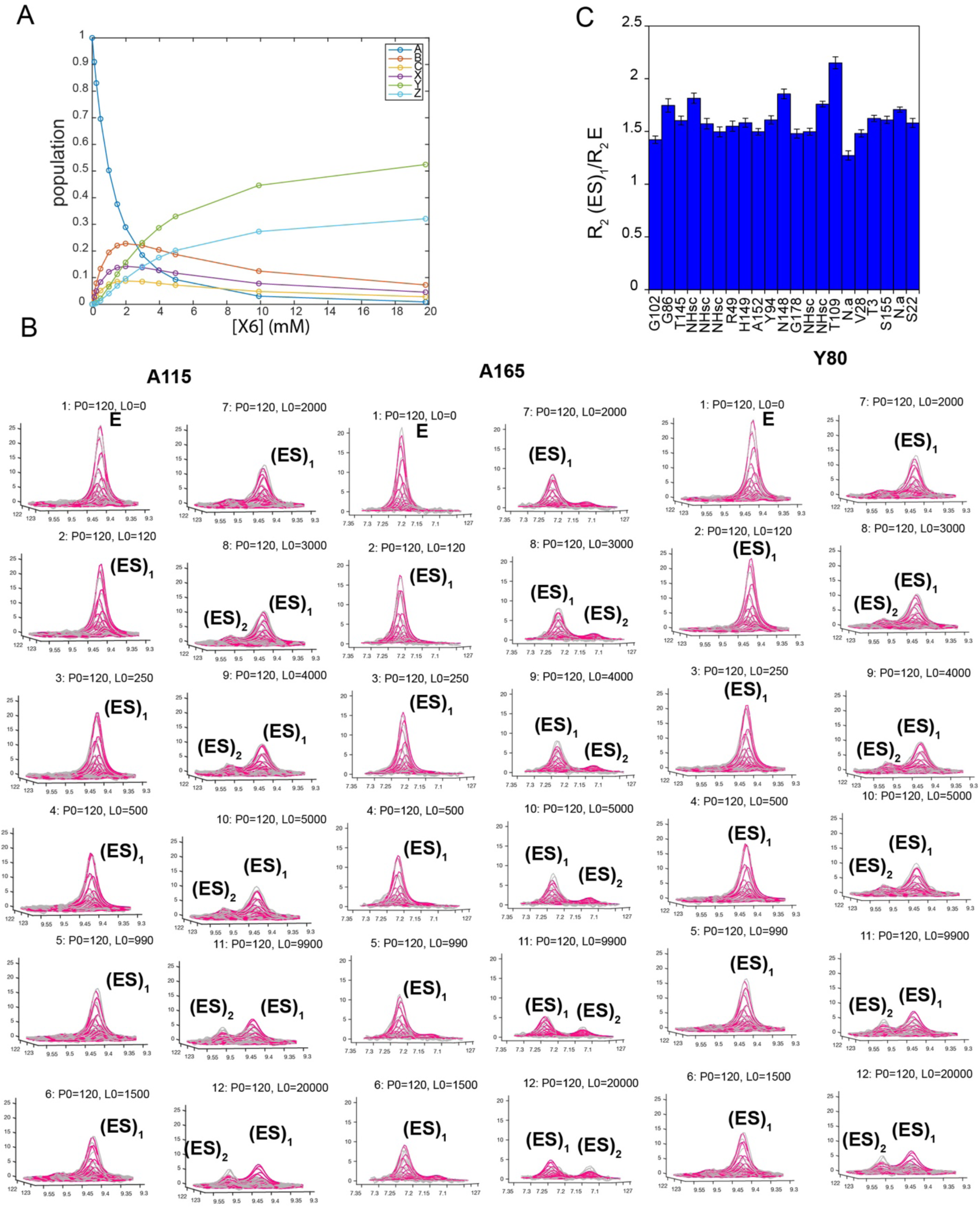
3D lineshape analysis of BCX E78Q titration with X6. (**A**) BCX-X6 bound states populations over the course of the titration, extracted from the global fit (see **M&M**). (**B**) Three representative examples of 3D peak shape fitting using TITAN of resonances observed in the BCX titration with X6. Red lines indicate the simulated data and grey the experimental. P0 and L0 are for the enzyme and ligand concentrations in µM, respectively. (**C**) General line broadening of the BCX E78Q upon titration with X6. The ratios between the R_2_ values determined from the 3D peaks shape analysis with TITAN in the absence and the presence of 20 mM X6 from amides that do not shift during titration. NHsc indicate side chain amides. N.a indicate non-assigned backbone amides.

**Fig. S4.**
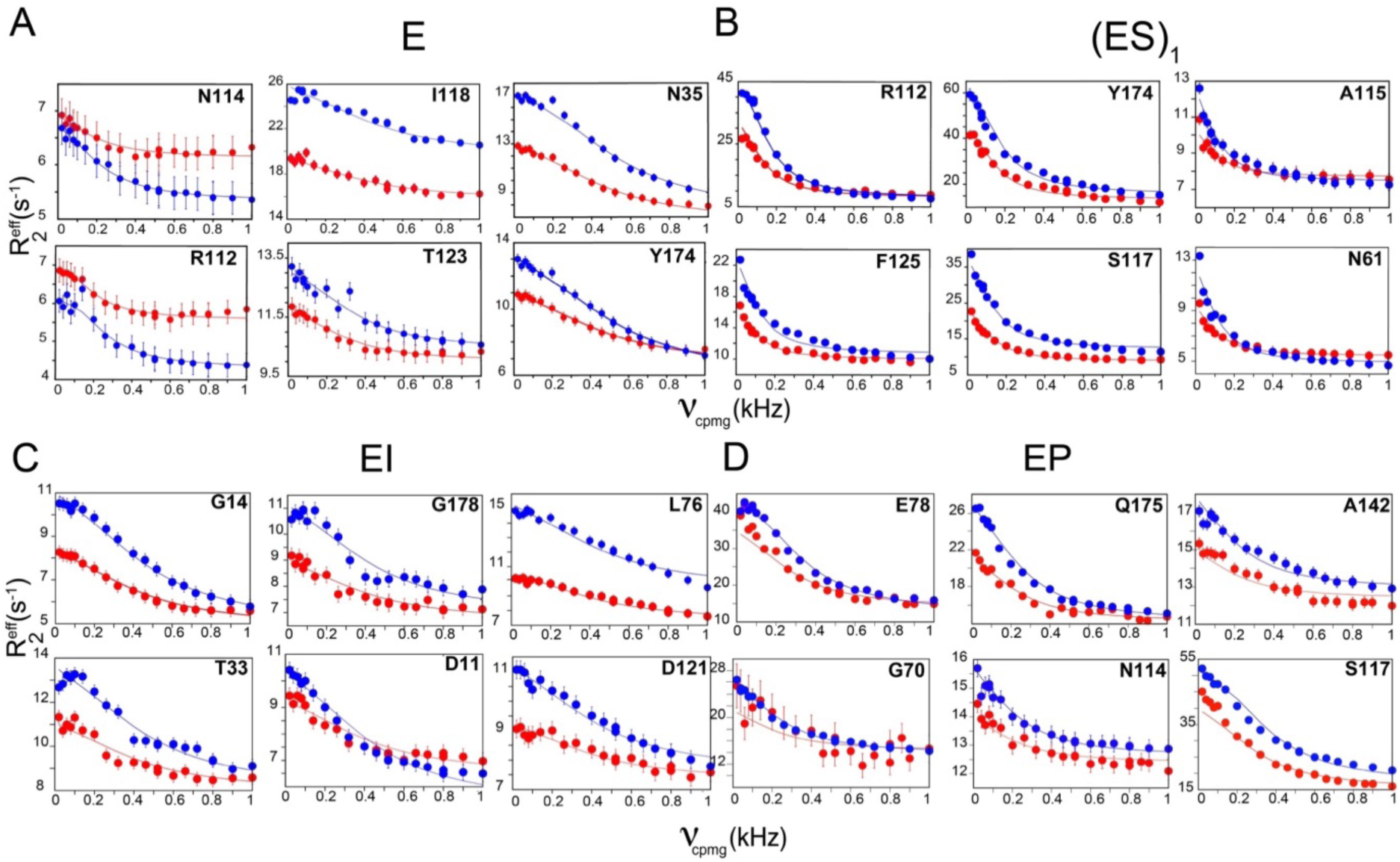
Global fit of CPMG RD data to a two-state chemical exchange model acquired on two static fields (red at 14 T and blue at 20 T) for (**A**) E; (**B**) (ES)_1_ (**C**) EI; (**D**) EP. The data of six representative residues of the global fit profile are shown for each state. Residue names are indicated.

**Fig. S5.**
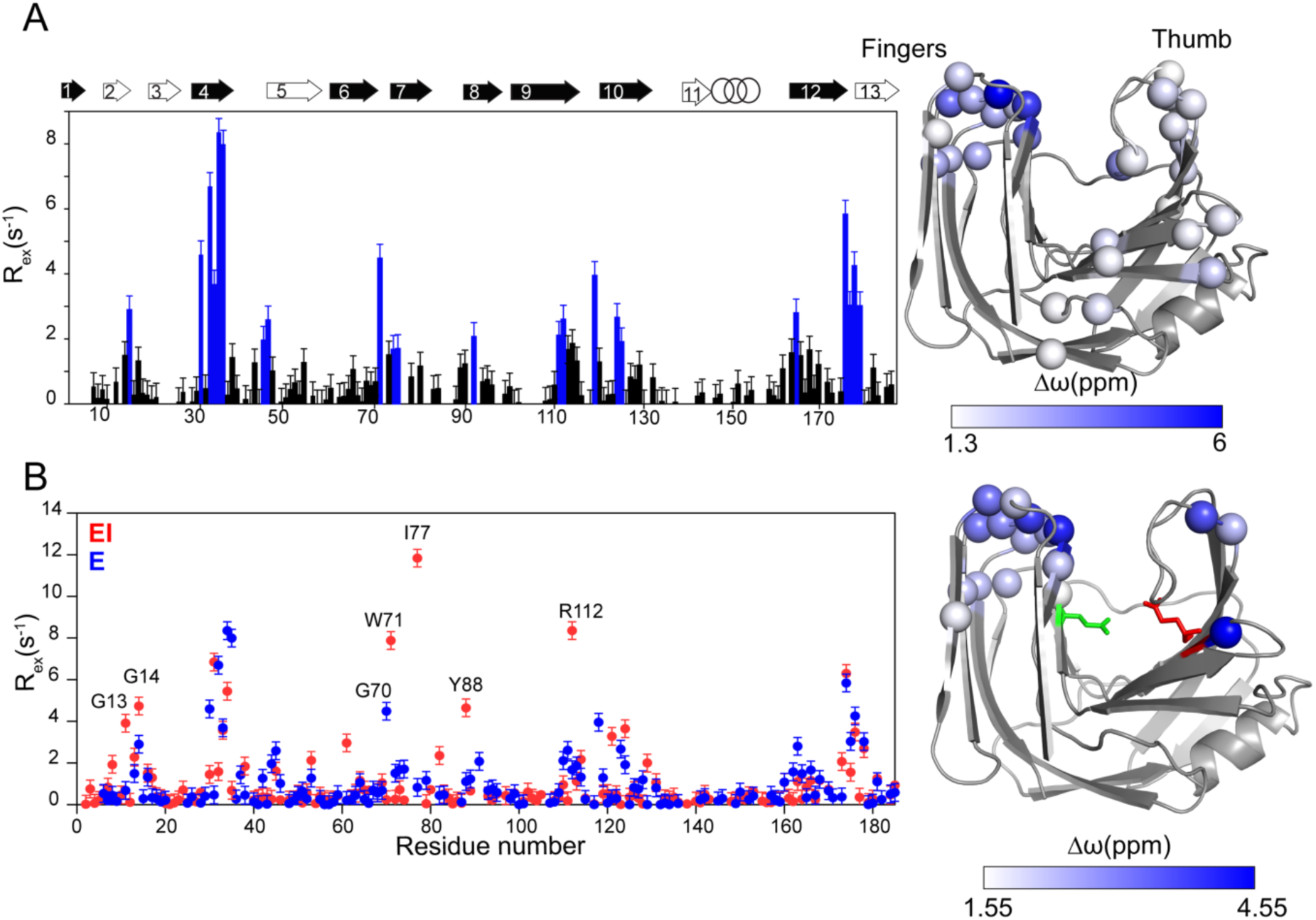
BCX milliseconds dynamics in the E and EI states. (**A**) Plot of R_ex_ vs. residue number of BCX resting state (E). Residues used in the CPMG RD global fit are shown in blue. The error bars represent the estimated error based on triplicate measurements at different *v*_CPMG_ frequencies in the same experiment. The secondary structure elements of BCX are represented by black arrows for β-strands of sheet A and in white ones for sheet B and the α-helix in rings. The “thumb” loop connects β-strands 9 and 10 ^12^. Amide ^15^N atoms used in the CPMG RD two-state global fit [k_ex_ = 2.39 (±0.07) × 10^3^ s^-1^; p_B_ = 0.8%] are shown in spheres on the BCX crystal structure and colored using a white/blue gradient by their predicted δω. (**B**) Overlay between the R_ex_ values of the EI (red dots) and the E states (blue dots) plotted against the residue number. Residues that show differences in their R_ex_ between the E and EI states are labelled. Amide ^15^N used in RD two-state global fitting [k_ex_ =2.05 (± 0.08) × 10^3^ s^-1^; p_B_ = 0.6%] are shown as spheres on the BCX crystal structure and colored by their predicted δω using a white/blue gradient. The red and green sticks indicate the nucleophile E78 and the acid/base E172 catalytic residues, respectively.

**Fig. S6.**
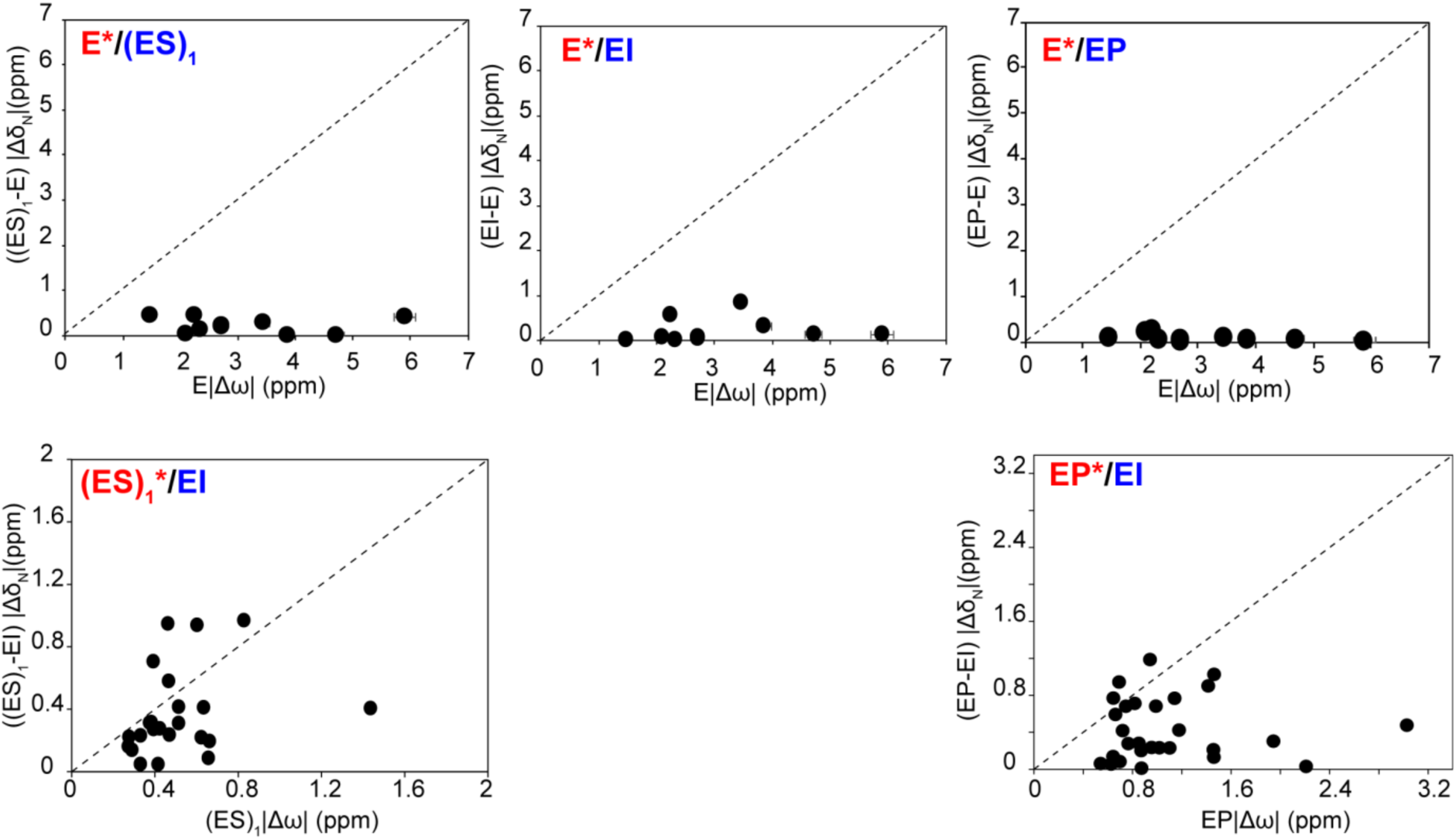
Correlation between the *Δω* from global fits of the RD data with the |*Δ*δ_N_| from the NMR titration data. At the top the *Δω* of the E excited state plotted against the |*Δ*δ _N_| of the (ES)_1_, EI and EP ground states. Below, the same is plotted for the (ES)_1_ (left) and EP (right) states vs. the CSP for the EI state. Excited states are indicated in red and marked with asterisks and ground states in blue.

**Fig. S7.**
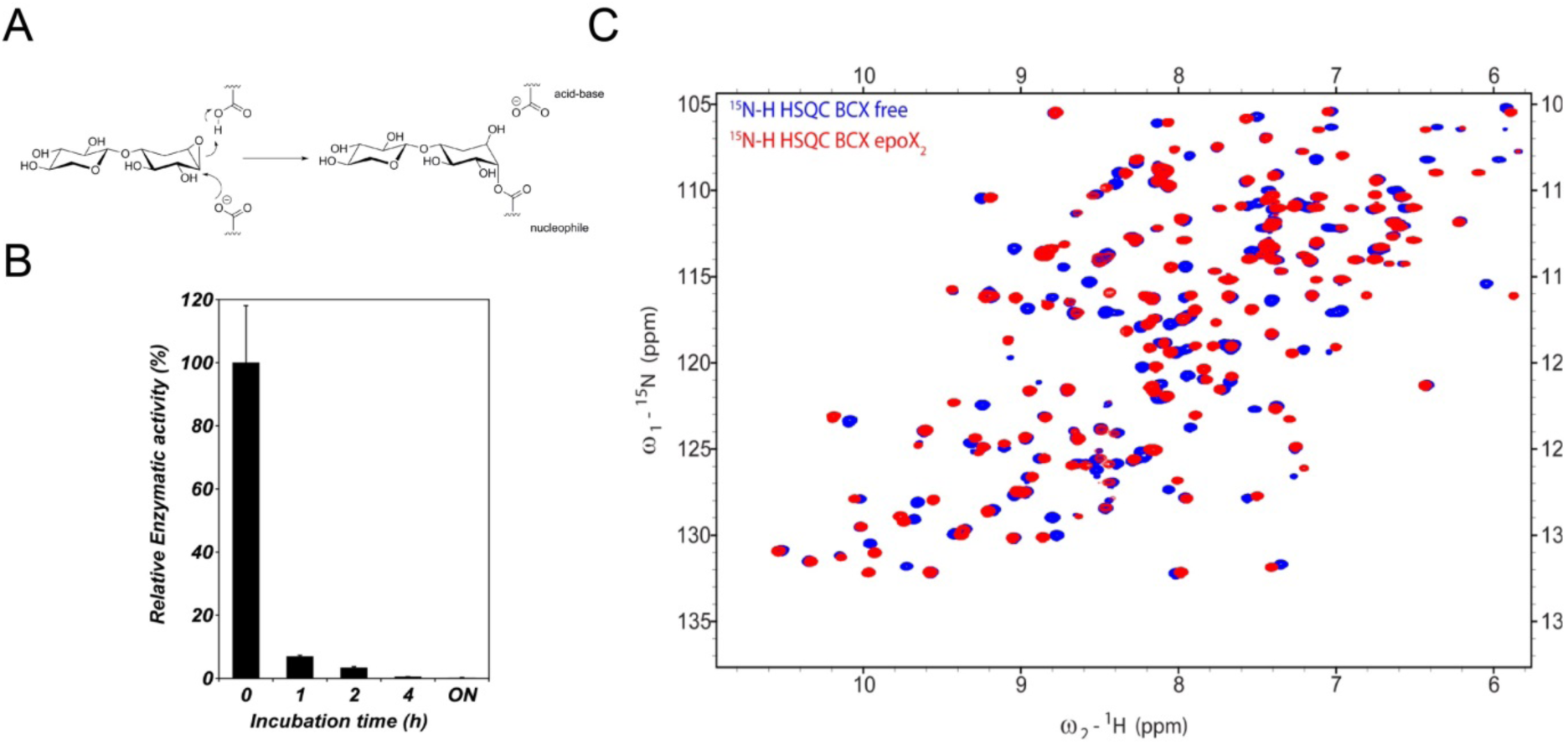
BCX EI complex. (**A**) BCX EI covalent complex formation by epoxyX2. The inactivation mechanism involves the attack of the ligand epoxide active center by the enzyme nucleophile, resulting in ring opening and the formation of a covalent bond between the enzyme nucleophile and the inhibitor, which emulates the EI state of the natural substrate hydrolysis reaction. The process is facilitated by protonation of the inactivator reactive center by the general acid/base residue (**B**) Residual BCX activity monitored over the course of incubation at 30°C in the presence of 1:20 (BCX: epoxyX_2_) ratio. Error bars indicate ±SD of a duplicate. (**C**) ^1^H-^15^N HSQC spectra overlay of the E and EI states.

**Fig. S8.**
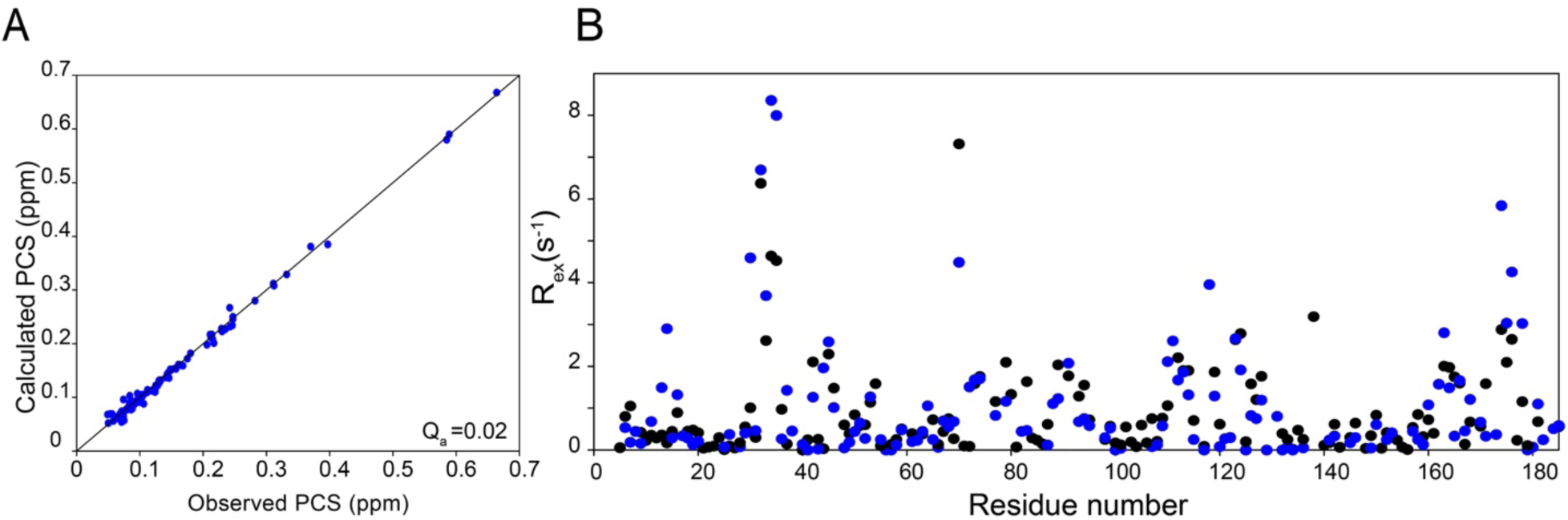
BCX E78Q conformation and milliseconds dynamics relaxation. (**A**) Correlation between the BCX E78Q observed vs. predicted PCSs from the fit to the BCX crystal structure (PDB ID:2bvv). (**B**) Overlay between the R_ex_ values of the BCX WT (blue) and the BCX E78Q (black).

**Table S1.**
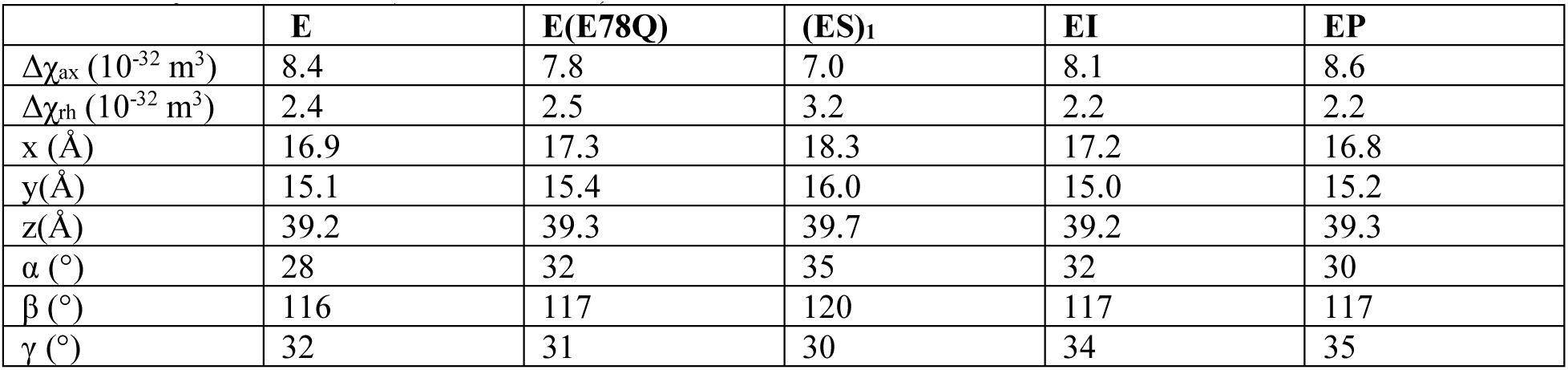
Tensor parameters of BCX in the different substrate bound states obtained from the PCS fit on the free enzyme structure (PDB ID:2bvv).

**Table S2.** Global fit parameters of BCX E78Q titration with X6 (see **M&M)**

| Parameter | value |
| --- | --- |
| $K_D \text{ AB}$ | $2.84 \pm 0.13 \text{ mM}$ |
| $K_D \text{ AX}$ | $3.4 \pm 0.1 \text{ mM}$ |
| $K_D \text{ CZ}$ | $1.7 \pm 0.3 \text{ mM}$ |
| $k_{off} \text{ AB}$ | $1122 \pm 284 \text{ s}^{-1}$ |
| $k_{off} \text{ AX}$ | $11489 \pm 807 \text{ s}^{-1}$ |
| $k_{ex} \text{ BC}$ | $71 \pm 5 \text{ s}^{-1}$ |
| $K_{BC}$ | 0.4 |
| $K_{YZ}$ | 0.8 |

**Table S3.**
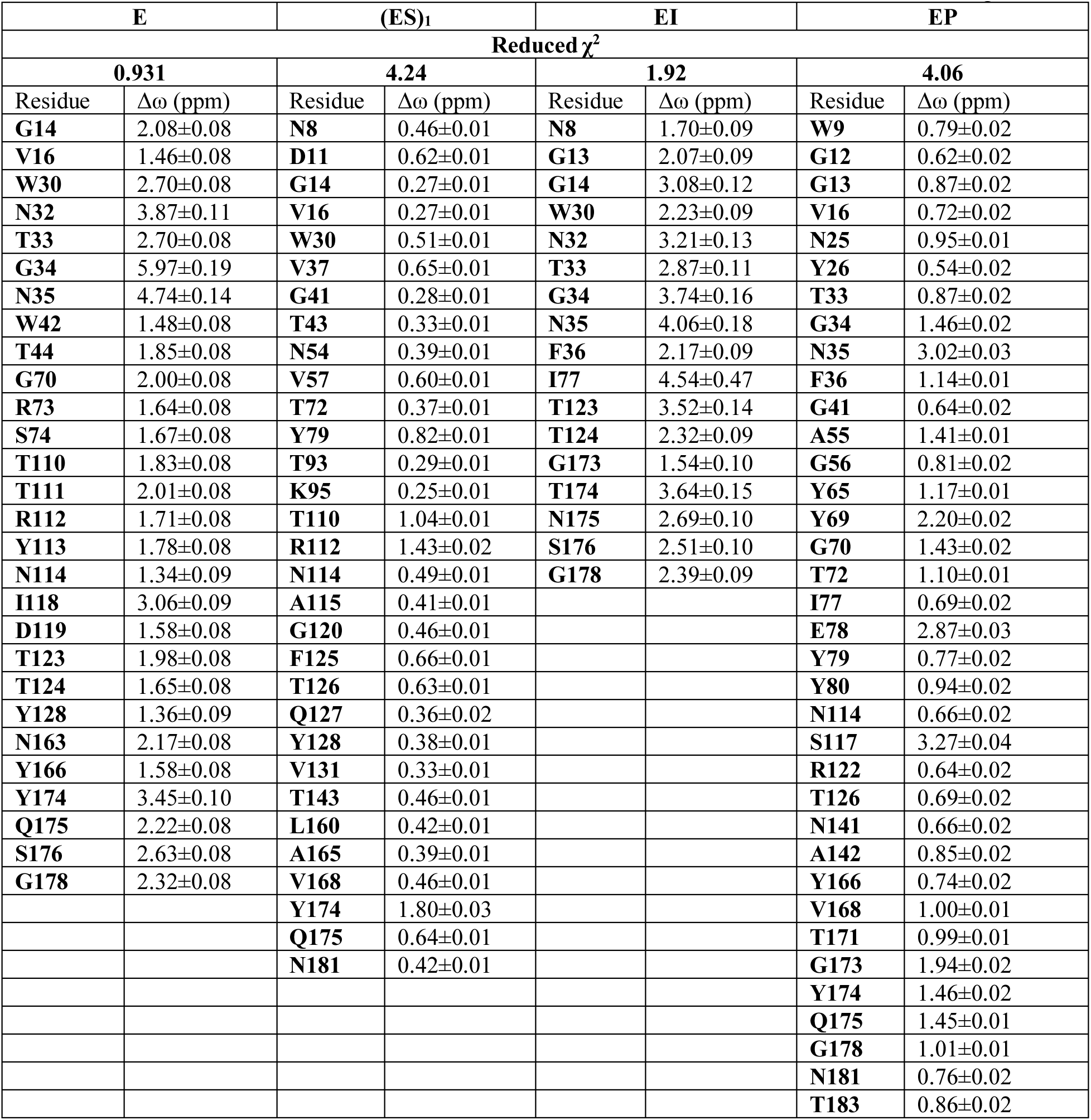
BCX CPMG-RD δω values for each substrate bound state obtained from a two-state global fit.

| E |  | (ES)<sub>1</sub> |  | EI |  | EP |  |
| --- | --- | --- | --- | --- | --- | --- | --- |
| Reduced $\chi^2$ | | | | | | | |
| 0.931 |  | 4.24 |  | 1.92 |  | 4.06 |  |
| Residue | $\Delta\omega$ (ppm) | Residue | $\Delta\omega$ (ppm) | Residue | $\Delta\omega$ (ppm) | Residue | $\Delta\omega$ (ppm) |
| G14 | 2.08±0.08 | N8 | 0.46±0.01 | N8 | 1.70±0.09 | W9 | 0.79±0.02 |
| V16 | 1.46±0.08 | D11 | 0.62±0.01 | G13 | 2.07±0.09 | G12 | 0.62±0.02 |
| W30 | 2.70±0.08 | G14 | 0.27±0.01 | G14 | 3.08±0.12 | G13 | 0.87±0.02 |
| N32 | 3.87±0.11 | V16 | 0.27±0.01 | W30 | 2.23±0.09 | V16 | 0.72±0.02 |
| T33 | 2.70±0.08 | W30 | 0.51±0.01 | N32 | 3.21±0.13 | N25 | 0.95±0.01 |
| G34 | 5.97±0.19 | V37 | 0.65±0.01 | T33 | 2.87±0.11 | Y26 | 0.54±0.02 |
| N35 | 4.74±0.14 | G41 | 0.28±0.01 | G34 | 3.74±0.16 | T33 | 0.87±0.02 |
| W42 | 1.48±0.08 | T43 | 0.33±0.01 | N35 | 4.06±0.18 | G34 | 1.46±0.02 |
| T44 | 1.85±0.08 | N54 | 0.39±0.01 | F36 | 2.17±0.09 | N35 | 3.02±0.03 |
| G70 | 2.00±0.08 | V57 | 0.60±0.01 | I77 | 4.54±0.47 | F36 | 1.14±0.01 |
| R73 | 1.64±0.08 | T72 | 0.37±0.01 | T123 | 3.52±0.14 | G41 | 0.64±0.02 |
| S74 | 1.67±0.08 | Y79 | 0.82±0.01 | T124 | 2.32±0.09 | A55 | 1.41±0.01 |
| T110 | 1.83±0.08 | T93 | 0.29±0.01 | G173 | 1.54±0.10 | G56 | 0.81±0.02 |
| T111 | 2.01±0.08 | K95 | 0.25±0.01 | T174 | 3.64±0.15 | Y65 | 1.17±0.01 |
| R112 | 1.71±0.08 | T110 | 1.04±0.01 | N175 | 2.69±0.10 | Y69 | 2.20±0.02 |
| Y113 | 1.78±0.08 | R112 | 1.43±0.02 | S176 | 2.51±0.10 | G70 | 1.43±0.02 |
| N114 | 1.34±0.09 | N114 | 0.49±0.01 | G178 | 2.39±0.09 | T72 | 1.10±0.01 |
| I118 | 3.06±0.09 | A115 | 0.41±0.01 |  |  | I77 | 0.69±0.02 |
| D119 | 1.58±0.08 | G120 | 0.46±0.01 |  |  | E78 | 2.87±0.03 |
| T123 | 1.98±0.08 | F125 | 0.66±0.01 |  |  | Y79 | 0.77±0.02 |
| T124 | 1.65±0.08 | T126 | 0.63±0.01 |  |  | Y80 | 0.94±0.02 |
| Y128 | 1.36±0.09 | Q127 | 0.36±0.02 |  |  | N114 | 0.66±0.02 |
| N163 | 2.17±0.08 | Y128 | 0.38±0.01 |  |  | S117 | 3.27±0.04 |
| Y166 | 1.58±0.08 | V131 | 0.33±0.01 |  |  | R122 | 0.64±0.02 |
| Y174 | 3.45±0.10 | T143 | 0.46±0.01 |  |  | T126 | 0.69±0.02 |
| Q175 | 2.22±0.08 | L160 | 0.42±0.01 |  |  | N141 | 0.66±0.02 |
| S176 | 2.63±0.08 | A165 | 0.39±0.01 |  |  | A142 | 0.85±0.02 |
| G178 | 2.32±0.08 | V168 | 0.46±0.01 |  |  | Y166 | 0.74±0.02 |
|  |  | Y174 | 1.80±0.03 |  |  | V168 | 1.00±0.01 |
|  |  | Q175 | 0.64±0.01 |  |  | T171 | 0.99±0.01 |
|  |  | N181 | 0.42±0.01 |  |  | G173 | 1.94±0.02 |
|  |  |  |  |  |  | Y174 | 1.46±0.02 |
|  |  |  |  |  |  | Q175 | 1.45±0.01 |
|  |  |  |  |  |  | G178 | 1.01±0.01 |
|  |  |  |  |  |  | N181 | 0.76±0.02 |
|  |  |  |  |  |  | T183 | 0.86±0.02 |

**Table S4.** Kinetic parameters. Microscopic rate constants (grey highlight) of BCX catalytic cycle of the model shown in **Figure 4** that are in agreement the observed kinetic parameters. k_ex_(a<>E) and k_ex_ (a<>b) refer to the exchange from the major (ES)_1_ state a to the free state E and to the minor state b via the free state E, respectively. Analogous for c and d for the EP state. The constants labelled ‘app’ are the experimentally observed apparent constants. Note that the microscopic rate constants merely serve to demonstrate that with reasonable values the experimental data can be reproduced.

|  | (ES)<sub>1</sub> |  | EP |
| --- | --- | --- | --- |
| $k_{\text{on},a}$ (ES)<sub>1</sub> (M<sup>-1</sup>s<sup>-1</sup>) | 3.6×10<sup>5</sup> | $k_{\text{on},c}$ (EP) (M<sup>-1</sup>s<sup>-1</sup>) | 3.0×10<sup>5</sup> |
| $k_{\text{on},b}$ (ES)<sub>1</sub>* (M<sup>-1</sup>s<sup>-1</sup>) | 2.3×10<sup>4</sup> | $k_{\text{on},d}$ (EP)* (M<sup>-1</sup>s<sup>-1</sup>) | 6.0×10<sup>3</sup> |
| $k_{\text{off},a}$ (ES)<sub>1</sub> (s<sup>-1</sup>) | 1160 | $k_{\text{off},c}$ (EP) (s<sup>-1</sup>) | 3150 |
| $k_{\text{off},b}$ (ES)<sub>1</sub>* (s<sup>-1</sup>) | 730 | $k_{\text{off},d}$ (EP)* (s<sup>-1</sup>) | 1500 |
| $k_{\text{ex}}(a \rightleftharpoons b)$ (s<sup>-1</sup>) | 741 (±9) | $k_{\text{ex}}(c \rightleftharpoons d)$ (s<sup>-1</sup>) | 1530 (±18) |
| pB | 9.2% (± 0.2) | pB | 4% |
| $k_{\text{ex}}^{\text{app}}(a \rightleftharpoons E)$ (s<sup>-1</sup>) | 0.9 (±0.25)×10<sup>4</sup> | $k_{\text{ex}}^{\text{app}}(c \rightleftharpoons E)$ (s<sup>-1</sup>) | 4.3 (±0.3)×10<sup>4</sup> |
| $k_{\text{off}}^{\text{app}}$ (s<sup>-1</sup>) | 1.1 ×10<sup>3</sup> (±0.3) | $k_{\text{off}}^{\text{app}}$ (s<sup>-1</sup>) | 3.1 (±0.2) ×10<sup>3</sup> |
| $K_d^{\text{app}}$ (mM) | 2.8 (±0.1) | $K_d^{\text{app}}$ (mM) | 10.00 (±0.1) |

